# Biophysical principles of choanoflagellate self-organization

**DOI:** 10.1101/659698

**Authors:** Ben T. Larson, Teresa Ruiz-Herrero, Stacey Lee, Sanjay Kumar, L. Mahadevan, Nicole King

**Affiliations:** Howard Hughes Medical Institute and the Department of Molecular and Cell Biology, University of California, Berkeley, CA, USA; Biophysics Graduate Group, University of California, Berkeley, CA, USA; Paulson School of Engineering and Applied Sciences, Harvard University, Cambridge, MA, USA; UC Berkeley-UCSF Graduate Program in Bioengineering, Department of Bioengineering, University of California, Berkeley, CA, USA; Department of Chemical and Biomolecular Engineering, University of California, Berkeley, CA, USA; Departments of Physics and Organismic and Evolutionary Biology, and Kavli Institute for NanoBio Science and Technology, Harvard University, Cambridge, MA, USA

**Keywords:** morphogenesis, multicellularity, quantitative microscopy, physical constraints, extracellular matrix, morphospace

## Abstract

Inspired by the patterns of multicellularity in choanoflagellates, the closest living relatives of animals, we quantify the biophysical processes underlying the morphogenesis of rosette colonies in the choanoflagellate *Salpingoeca rosetta*. We find that rosettes reproducibly transition from an early stage of 2D growth to a later stage of 3D growth, despite the underlying stochasticity of the cell lineages. We postulate that the extracellular matrix (ECM) exerts a physical constraint on the packing of proliferating cells, thereby sculpting rosette morphogenesis. Our perturbative experiments coupled with biophysical simulations demonstrates the fundamental importance of a basally-secreted ECM for rosette morphogenesis. In addition, this yields a morphospace for the shapes of these multicellular colonies, consistent with observations of a range of choanoflagellates. Overall, our biophysical perspective on rosette development complements previous genetic perspectives and thus helps illuminate the interplay between cell biology and physics in regulating morphogenesis.

**Significance statement:** Comparisons among animals and their closest living relatives, the choanoflagellates, have begun to shed light on the origin of animal multicellularity and development. Here we complement previous genetic perspectives on this process by focusing on the biophysical principles underlying colony morphology and morphogenesis. Our study reveals the crucial role of the extracellular matrix in shaping the colonies and leads to a phase diagram that delineates the range of morphologies as a function of the biophysical mechanisms at play.

## Introduction

Nearly all animals start life as a single cell (the zygote) that, through cell division, cell differentiation, and morphogenesis, gives rise to a complex multicellular adult form (1, 2). These processes in animals require regulated interplay between active cellular processes and physical constraints (3–9). A particularly interesting system in which to study this interplay is the choanoflagellates, the closest relatives of animals (10–12). Choanoflagellates are aquatic microbial eukaryotes whose cells bear a diagnostic “collar complex” composed of an apical flagellum surrounded by an actin-filled collar of microvilli (13, 14) (Fig. 1). The life histories of many choanoflagellates involve transient differentiation into diverse cell types and morphologies (15, 16). For example, in the model choanoflagellate *Salpingoeca rosetta*, solitary cells develop into multicellular colonies through serial rounds of cell division (17), akin to the process by which animal embryos develop from a zygote (Fig. 1A). Therefore, choanoflagellate colony morphogenesis presents a simple, phylogenetically-relevant system for investigating multicellular morphogenesis from both a biological and a physical perspective (14).

**Figure 1.**
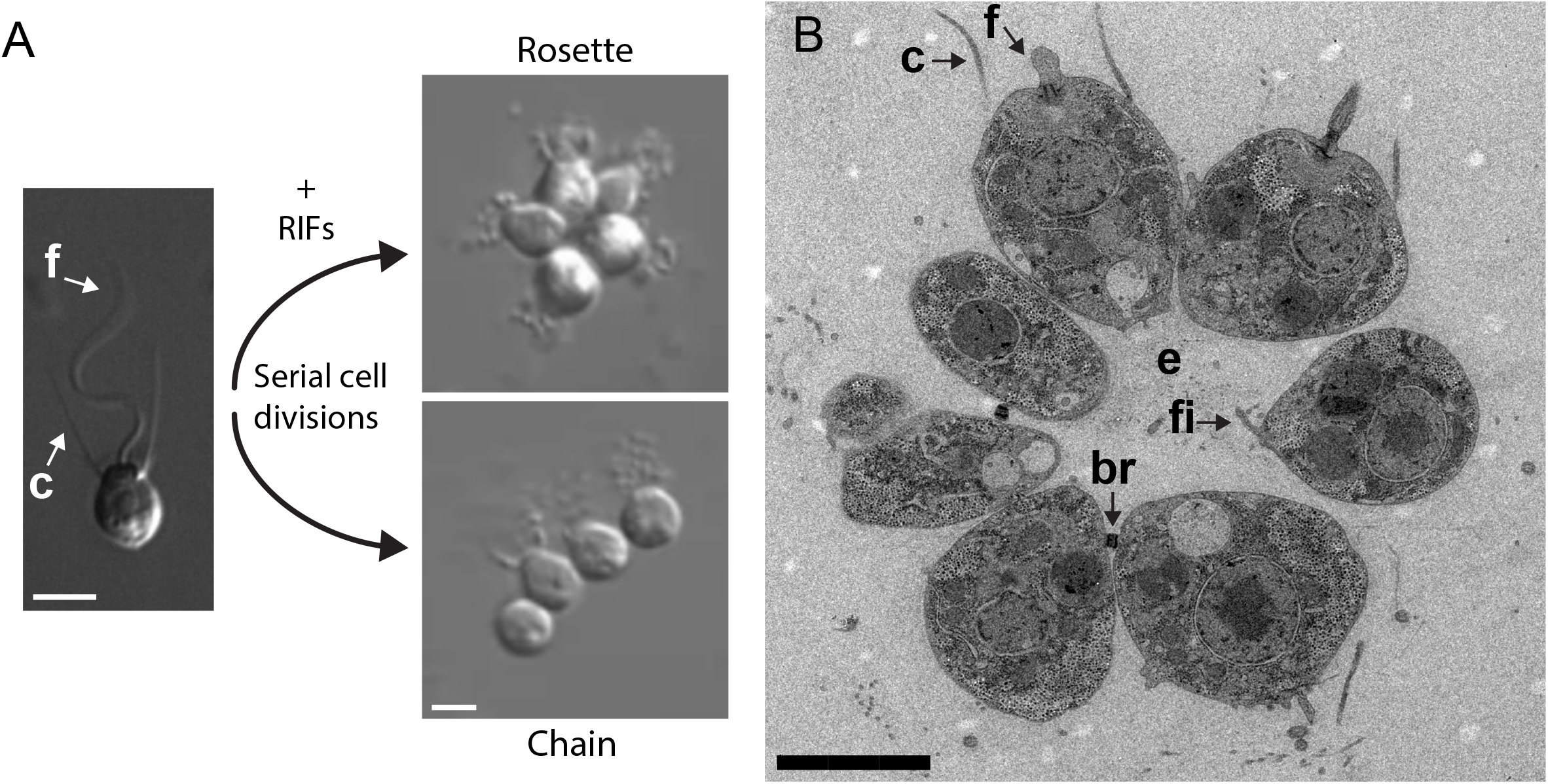
The choanoflagellate *Salpingoeca rosetta* develops from a single cell into multicellular colonies through serial rounds of cell divisions (17). **(A)** All choanoflagellate cells bear a diagnostic “collar complex” composed of an apical flagellum (f) surrounded by an actin-filled collar of microvilli (c) (13, 14). *S. rosetta* produces two different colonial forms depending on environmental conditions: compact, mechanically robust, roughly spherical rosette colonies (Rosette) that form in the presence of specific bacterially produced Rosette Inducing Factors (RIFs; (16, 17, 23, 24)), and fragile, linear chain colonies (Chain) that form during rapid cell growth in the absence of RIFs (16). Both types of colonies form developmentally by serial cell divisions. Single cell image adapted from (16), and rosette and chain images adapted from (19). **(B)** A thin section through the equator of a rosette, imaged by transmission electron microscopy, reveals the subcellular architecture of a rosette. Cells in rosettes are packed close to one another around a central focus with the collar complex of each cell facing outward into the environment (c=collar and f=flagellum). Most cells are connected to one another by thin cytoplasmic bridges (br, only two of which are visible in this section) (37), which are also present in chains (16). The center of rosettes is devoid of cells but is filled with a secreted extracellular matrix (e, faintly visible here as granular material), into which cells extend filopodia (fi) (16, 18). All scale bars = 3 μm.

*S. rosetta* forms planktonic rosette-shaped colonies (“rosettes”), in which the cells are tightly packed into a rough sphere that resembles a morula-stage animal embryo (17). Because the cell division furrow forms along the apical-basal axis, thereby dissecting the collar, all of the cells in rosettes are oriented with their flagella and collars facing out into the environment and their basal poles facing into the rosette interior (Fig. 1B). Interestingly, all three genes known to be required for rosette development are regulators of the extracellular matrix (ECM): a C-type lectin called *rosetteless* (18) and two predicted glycosyltransferases called *jumble* and *couscous* (19). Nonetheless, little is known about either the mechanistic role of the ECM or the extent to which rosette morphogenesis is shaped by physical constraints.

A critical barrier to understanding the biological and physical mechanisms underlying rosette morphogenesis has been the absence of a detailed characterization of the morphogenetic process. For example, it is not known whether rosettes form through the development of invariant cell lineages akin to those seen in *C. elegans* (20) or through stochastic cell divisions, as occurs, for example, in sponges and mice (21, 22). Moreover, it is not known whether there are identifiable developmental stages in rosette development. To quantify the principles of rosette morphogenesis, we used a combination of quantitative descriptions of rosette development, experimental perturbations, and biophysical simulations that together reveal the importance of the regulated secretion of basal ECM in physically constraining proliferating cells and thereby sculpting choanoflagellate multicellularity.

## Results

### Rosette morphogenesis displays a stereotyped transition from 2D to 3D growth

To constrain our search for mechanistic principles, we first quantified the range of sizes and spectrum of morphologies of *S. rosetta* rosettes by measuring the population-wide distribution of rosette size in terms of cell number. *S. rosetta* cultured solely in the presence of the rosette-inducing bacterium *A. machipongonensis* (23), lead to a population with a stationary cell number distribution. While some rosettes contained as many as 25 cells, the most common rosette size was 8 cells/rosette, with 51% of rosettes containing between 6-8 cells (Fig. 2A). While rosettes grow through cell division, their ultimate size is determined by either colony fission (as previously reported; 16) or cell extrusion (Fig S1). In each case, the rosettes contained 8 or more cells, suggesting that these rosette size decreasing phenomena are more common in larger rosettes.

**Figure 2.**
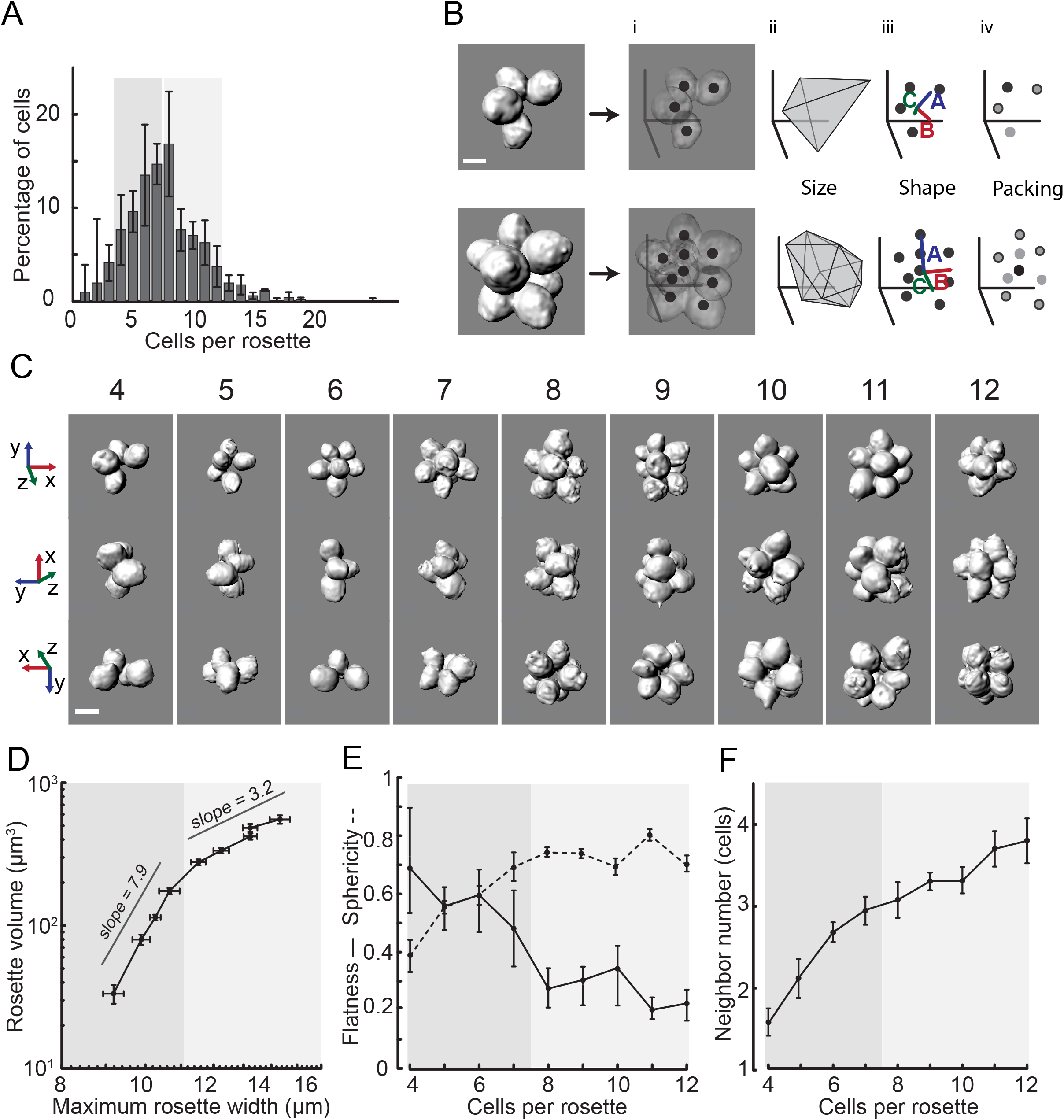
Quantitative analysis of rosette morphology reveals that rosettes undergo a reproducible 2D-3D growth transition. **(A)** In cultures grown under conditions of constant rosette induction, *S. rosetta* existed as unicells, doublets, triplets, and rosettes containing between 4 - 25 cells (an “individual” refers to any unicell or group of cells in each of these categories), with the most common rosette size being 8 cells/individual. Shown is the mean percentage of total cells in a population (y-axis) found in single cells, cell doublets, cell triplets, and rosettes of increasing size, plotted by number of cells/rosette (x-axis). Error bars indicate standard deviations from measurements obtained on three different days. N = 511. **(B)** Our image analysis pipeline allowed us to quantify and compare rosette morphology and is illustrated here for two representative rosettes. From left to right, for each rosette, (i) cell positions were extracted from segmented images and then used to determine aspects of rosette morphology including (ii) rosette size, including volume (measured by generating a convex hull), (iii) shape, including flatness and sphericity (the former quantified by 1 − *C*/*B* and the latter by 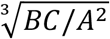 where *A, B*, and *C* are the principle axes in descending order by magnitude of a principle components analysis-based ellipsoid fit of cell positions), (iv) and cell packing (neighbor number determined by Voronoi tessellation(80)). **(C)** Representative rosettes are shown in three roughly orthogonal views for size classes ranging from four to 12 cells/rosette, with the numbers above each image column indicating the number of cells/rosette. Following previous work (24), we defined four cells as the smallest number of cells clearly identifiable as a rosette. **(D)** Rosettes transition from an early phase of major shape change (dark grey; scaling exponent ~8) to a later phase of approximately isotropic growth (light grey; scaling exponent ~3), as shown by a log-log plot of rosette volume (y-axis) vs. maximum rosette width (x-axis). **(E)** Rosettes transition from a relatively flat morphology during the 4-6 cell stage (dark grey; mean flatness ≅ 0.5 - 0.7 and mean sphericity ≅ 0.4 - 0.6, with flatness = 1.0 perfectly flat and sphericity = 1.0 perfectly spherical) to a more spheroidal morphology during the 8-12 cell stage (light grey; mean flatness ≅ 0.2 - 0.3 and mean sphericity ≅ 0.7-0.8). **(F)** Packing increases with number of cells at a decreasing rate. Points on plots D-F represent mean values; error bars indicate standard error of the mean. N = 100 rosettes, with at least 8 rosettes from each cell-number class, pooled from three different samples. All scale bars = 3 μm.

We next quantified defining features of the 3D morphology of rosettes containing between four (following (24), we defined four cells as the smallest cell number clearly identifiable as a rosette) to twelve cells (representing 90% of rosettes at steady state, Methods; Fig. 2B, C). This analysis revealed that rosettes increased in volume and diameter as cell number increased (Fig. 2D). Although the average cell volume reduced between the four-cell and five-cell stages of rosette development, average cell volume did not change substantially with increasing cell number after the five-cell stage (Fig. S2), suggesting that cells in rosettes grow between cell divisions. This contrasts with cleavage in the earliest stages of animal embryogenesis, in which cell volume steadily decreases as cell divisions proceed with no cell or overall tissue growth (2).

Our analyses revealed that rosette morphogenesis displays two distinct, but previously undescribed phases: (1) a 2D phase of growth from four to seven cells, during which the overall shape of rosettes changed substantially with increasing cell number and (2) a 3D phase from eight to twelve cells, during which rosettes expanded nearly isotropically (Fig. 2C-E). Interestingly, the most common rosette size (8 cells) corresponded to the transition between the two phases of growth.

Transitions from 2D to 3D growth can be driven by the constrained growth of cell layers leading to increasing mechanical stresses (25–29). We hypothesized that the physical packing of cells in rosettes might constrain cell growth and proliferation and help explain the growth transition during rosette morphogenesis. Indeed, cell packing initially increased (as indicated by an increase in the number of nearest neighbor cells, Fig. 2F, and suggested by the reduced average sphericity of cells, Fig. S2). Following the growth transition at the 8-cell stage, cell packing continued to increase with increasing cells/rosette, although the rate of increase slowed as a function of the number of cells/rosette (Fig. 2F). Therefore, the transition to isotropic 3D growth in eight-cell rosettes may occur in response to the accumulation of stress caused by the increase in cell packing in growing rosettes.

### Rosette developmental dynamics are stochastic

The influence of cell packing on rosette morphogenesis did not preclude the possibility that the rosette developmental program might also involve specific patterns of cell division that result in well-defined cell lineages. We therefore documented cell lineages in live, developing rosettes (Fig. 3). Consistent with the single previous published observation of live rosette development (17), the cells maintained polarity throughout development, with their division planes oriented along the apical-basal axis. Relative to the cell division times in linear chains (Fig. 1A), which form when rosette inducing bacteria are absent, we observed a slight but statistically significant increase in division rate in rosettes (p=0.03 by Wilcoxon rank sum test Fig. S3). In addition, we found that both the order and timing of cell divisions differed among different rosettes (Fig. 3B, C), ruling out the possibility that cell lineages are invariant. This process of apparently unpatterned cell divisions resembles the dynamics of early embryogenesis in diverse animals, including sponges and mice (21, 22).

**Figure 3.**
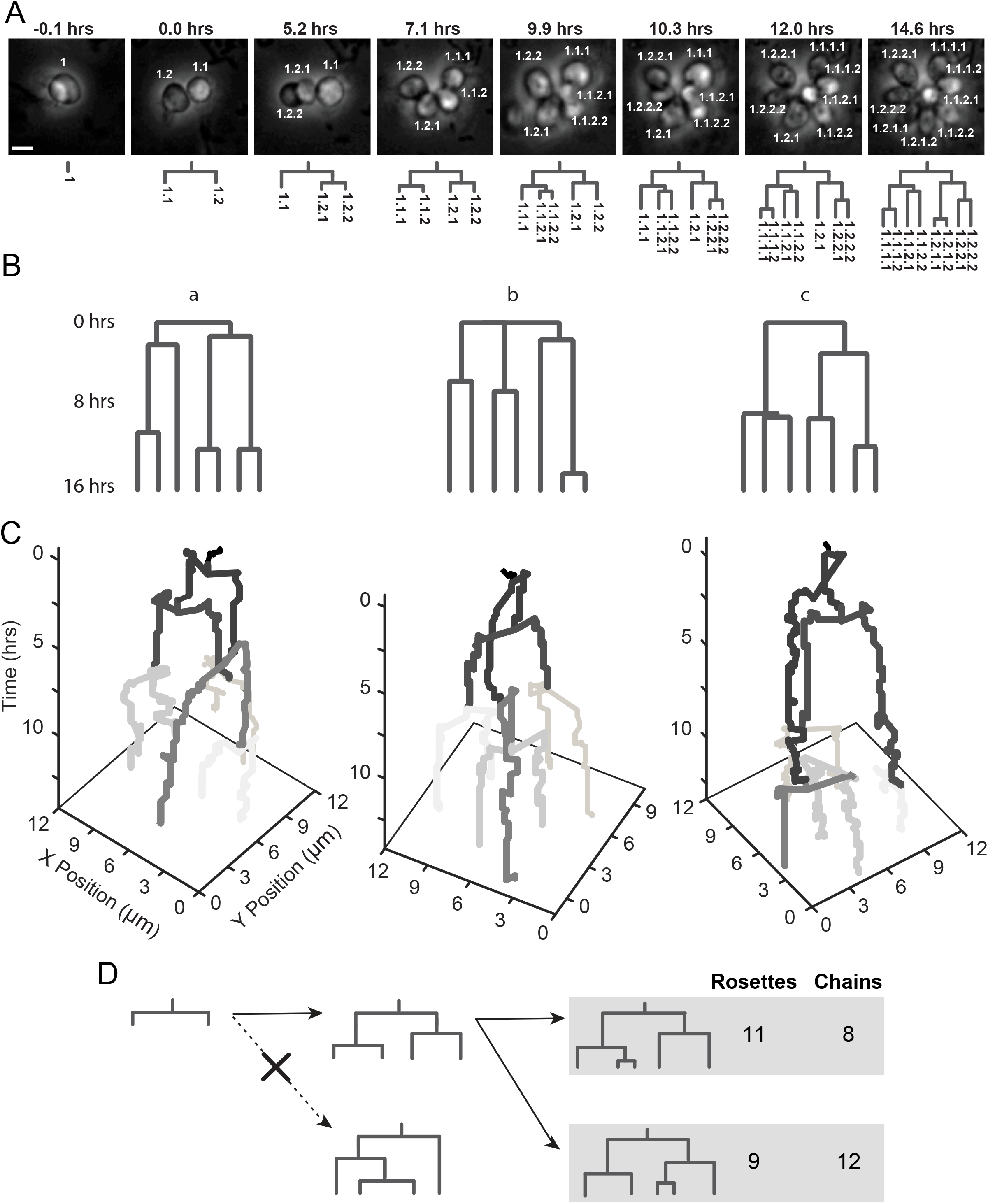
Stochasticity of developmental dynamics revealed by lineage analysis. **(A)** Lineage analysis for a representative rosette, imaged live by time lapse microscopy. Individual daughter cells were marked to record their relationship to their parent lineage (e.g. 1.1 and 1.2 are daughters of cell 1). Scale bar is 3 μm (B) Representative cell lineages during rosette development illustrating differences in both the order and timing of cell divisions, with the first two on the left having the same division order but with large differences in division times, and the lineage on the right showing differences in both division order and timing. Branch lengths scale with time and are set to zero based on the first division. **(C)** Lineages a-c from (B) displayed as space-time plots illustrate cell division variability between rosettes in both space and time. Plots also demonstrate that cells remain in place after divisions, with no large rearrangements, moving apart only slightly as they grow. Colors from dark to light gray indicate the order of cell divisions. **(D)** Cell lineages that form during rosette and chain development are balanced, although the specific times of cell divisions in different lineages and different rosettes or chains can differ. In rosettes and chains, imbalanced lineage structures (i.e. with significantly different numbers of cells) were not observed at any stage. Shown here are results for four and five-cell rosettes and chains. The dashed line with an “x” indicates that this division pattern was never observed. Furthermore, this uneven branching pattern was never observed in sub-lineages of any developing chain or rosette. Data were pooled from three different rosette induction experiments (for rosettes) and three different experiments with uninduced cells (for chains).

Although division patterns were variable between rosettes, ruling out the possibility of invariant cell lineages, in no rosette did cells from the first, second, or third cell division give rise to more than 60% of cells (Fig. 3D). Moreover, cell division remained balanced throughout rosette morphogenesis, with no cell lineage coming to dominate. Importantly, the cell lineages of chains showed the same kind of stochasticity and variability as rosettes (Fig. 3D). These observations suggest that rosette morphogenesis does not require the strongest forms of cell cycle control or coordination (i.e. the synchronous divisions or deterministic division timing or order observed in the development of some animals such as *C. elegans, Xenopus, Drosophila*, and zebrafish (30–33) and in the green alga *Volvox* (34–36)).

### ECM constrains proliferating cells in rosettes

To reconcile the stereotyped 3D growth transition (Fig. 2) with the stochastic developmental dynamics of rosette formation (Fig. 3), we set out to test the “ECM constraint hypothesis (Fig 4A, B).” This hypothesis was motivated by the idea that physical constraints imposed by the geometry and mechanics of cell packing play a key role in morphogenesis and that the source of the physical constraint in growing rosettes is the ECM, which is known to be required for rosette morphogenesis and connects all cells in a rosette, filling the rosette center (16, 18, 19, 37). The phenomenon of physically constrained morphogenesis suggests that the amount of ECM secreted during rosette development is an important factor in sculpting rosette morphogenesis (Fig. 4A, B). We visualized and quantified the volume of the ECM by staining with fluorescein-conjugated Jacalin, a galactose-binding lectin (19, 38). Importantly, Jacalin does not stain chains, so its target is likely specific to rosette ECM (19). We found that the relative amount of space occupied by basal ECM (ECM volume/total cell volume, denoted by ϕ) in developing rosettes was constant and maintained at roughly 6% (Fig. 4C). Therefore, we infer that cells in rosettes produce ECM at a constant rate relative to the growth of cells, either through synthesis and secretion alone or through a balance of regulated synthesis, secretion, and degradation.

**Figure 4.**
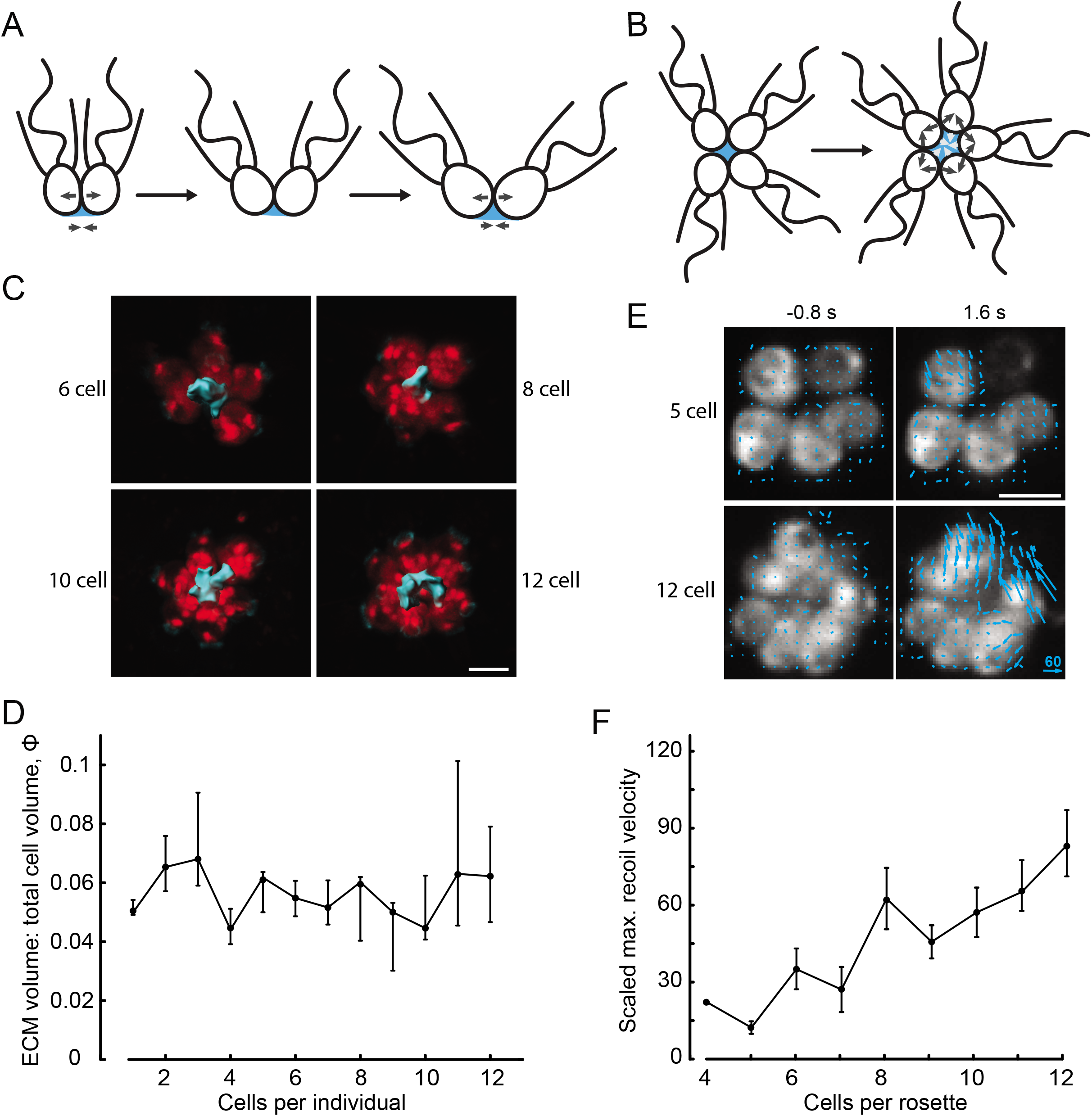
Extracellular matrix constrains proliferating cells during rosette morphogenesis. **(A-B)** Cartoons depicting ECM constraint hypothesis. **(A)** As cells grow, compressive forces exerted on neighboring cells (top set of arrows) may be balanced by stress in the basally-secreted ECM (blue), which resists deformation (bottom set of arrows). **(B)** If ECM is limiting, rosettes may undergo a jamming-like transition and accumulate residual stress as cells continue to grow and divide. **(C)** Representative images of 4 different rosettes of various sizes (of 6, 7, 8, and 10 cells/rosette ordered from left to right, top to bottom) with ECM tagged using Jacalin. **(D)** Ratio of total ECM volume to total cell volume as a function of cells/individual is maintained at a constant level over the course of rosette development. Points represent mean values, and error bars are bootstrap 95% confidence intervals. Data were collected from 93 rosettes pooled from two experiments, with at least 5 rosettes from each size class. Image processing and analysis proceeded similarly to that described in Fig. 1. **(E)** Laser ablation revealed increasing stress as a function of cells/rosette. Single cells in rosettes were ablated, and recoil velocities were measured by particle image velocimetry (PIV). Arrows in the images indicate the direction and magnitude of the velocity of the recoil as determined by PIV. Rosettes were found to always close when recoil was observed (in all cases beyond the 5-cell stage). **(F)** Recoil velocities (rescaled by the length scale of the average cell diameter of 5 μm and the time scale of the average division time of 6 hrs.) increased with increasing cells/rosette, indicating increasing stress as a function of cells/rosette (7, 39, 43). These results are consistent with the ECM constraint hypothesis and inconsistent with cytoplasmic bridges or cell-cell adhesion as the dominant factors stabilizing rosette structure. Points indicate mean values; error bars indicate standard error of the mean. Data were collected from 47 rosettes pooled from 3 different rosette inductions, with at least 4 rosettes from each size class. Scale bars = 5 μm.

A key prediction of the ECM constraint hypothesis (Fig. 4A, B) is that compressive stress on cells, balanced by stress in the ECM, should increase with cell number. Alternatively, cell-cell connections mediated by lateral cell-cell adhesion or cytoplasmic bridges formed during incomplete cytokinesis (16, 37) (Fig. 1B) might be primarily responsible for the structural integrity of rosettes. If cell-cell connections dominate over ECM in holding together rosettes, we would expect cells to be under tension such that measured stresses would be in the opposite direction to those predicted by the ECM constraint hypothesis.

To probe the balance of forces in developing rosettes, we performed laser ablation experiments, which provided a readout of the relative magnitude and direction of stresses within rosettes (39–42). Upon ablation of a single cell in a rosette, we found that the remaining cells immediately became more rounded and moved closer together, reducing the size of the gap left by the ablated cell (Fig. 4E. This result demonstrated that residual elastic stress (as measured by initial recoil velocity after ablation, (39)) is maintained in rosettes, with cells under compressive stress balanced by an additional component of residual stress. If rosettes were primarily held together by strong cell-cell adhesion or constrained by cytoplasmic bridges (Fig. 1B), the expected recoil would have been in the opposite direction, causing a larger gap to open in rosettes, due to cells increasing contact area with remaining neighbors in the former case and tension in bridges in the latter. Moreover, as the number of cells in rosettes increased, the measured residual stress increased (Fig. 4F), consistent with the ECM constraint hypothesis (Fig. 4A, B). These results ruled out strong cell-cell adhesion or constraint by cytoplasmic bridges as the dominant physical mechanisms underlying rosette integrity and morphogenesis.

Additionally, residual stress (as measured by initial recoil velocity (7, 39, 43)) displayed a sharp increase, by nearly a factor of two, at the 8-cell stage (Fig. 4F), coinciding with the 3D growth transition (Fig. 2). In conjunction with the observed increase in cell packing (Fig. 2F), this result suggested that the packing of cells is mechanically constrained in developing rosettes such that cells are increasingly compressed against one another with increasing cell number. We reasoned that the shared ECM secreted from the basal end of cells, adhesion to which is likely essential for rosette formation (18, 19), might be the source of this constraint. While we have ruled out bridges as a dominant component of the structural integrity of rosettes, they could play a role in stabilizing cell orientation to hinder out of plane growth during the 2D phase of rosette morphogenesis.

### Material properties of ECM affect morphogenesis

We next sought to test the ECM constraint hypothesis through perturbative experiments. While the hypothesis entails that changing geometrical properties such as cell shape and relative amount of ECM should have a substantial effect on rosette morphogenesis, these properties could not be experimentally tuned. However, we could perturb the mechanical properties of the ECM. To do so, we treated developing rosettes with strontium chloride (SrCl_2_). Strontium is a divalent cation that can stiffen hydrogels, including animal ECM, by increasing crosslinking density (44–48). Importantly, we found that SrCl_2_ has no detectable effect on cell growth at up to twice the highest concentration used during this set of experiments (Fig. S4). Under our ECM constraint hypothesis, we predicted that increased ECM stiffness would alter morphogenesis by further constraining cell packing, thus holding cells in a more compact arrangement along with a relative increase in residual stress. Consistent with our hypothesis, we found that rosettes became more compact with increasing SrCl_2_ concentration (Fig. 5A, B), and the 3D transition shifted to lower cell numbers, occurring at the five-cell stage for the highest SrCl_2_ concentration (Fig. 5C). Additionally, the transition to isotropic growth at the 8-cell stage was abolished (Fig. 5B). Together, these analyses reveal that morphogenesis is altered.

**Figure 5.**
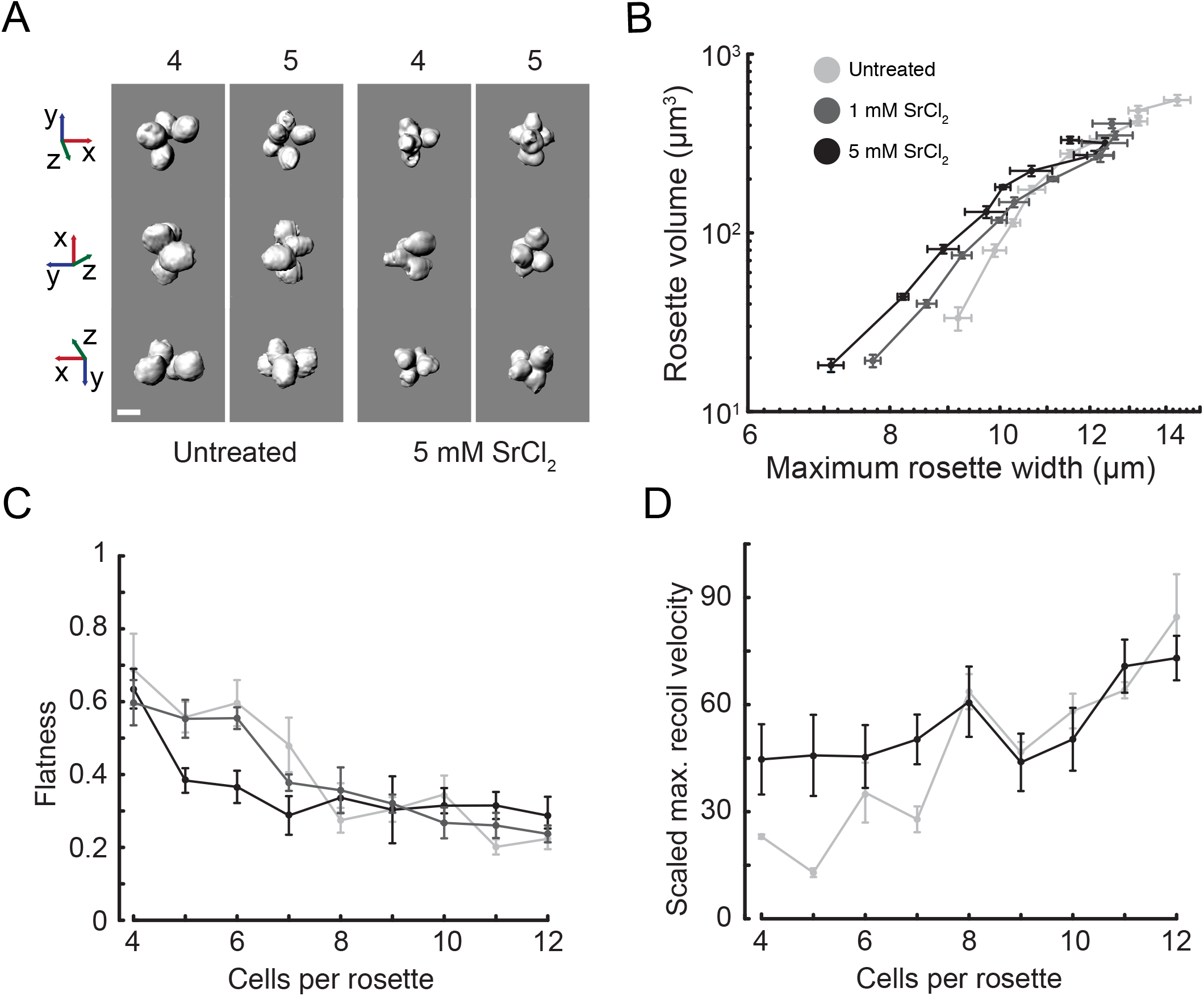
Material properties of the ECM affect morphogenesis, consistent with predictions of the ECM constraint hypothesis. Treatment of rosettes with SrCl_2_, which stiffens hydrogels by increasing crosslinking density (44–48), alters rosette morphogenesis. **(A)** Representative images of untreated and SrCl_2_-treated rosettes illustrate the change in rosette morphology. Cells were packed more tightly in SrCl_2_ treated rosettes, leading to differences in rosette size and shape. Scale bar = 4 μm. (B) The scaling relationship between maximum rosette width and volume revealed that SrCl_2_ abolished the transition to approximately isotropic growth observed in untreated rosettes. As in Fig. 2D, this is a log-log plot of rosette volume vs. maximum rosette width, with each point representing average values for a rosette cell-number class from 4-12 cells/rosette. Error bars are standard error of the mean. This analysis also revealed that rosettes became increasingly compact with increasing SrCl_2_ concentration. **(C)** Quantification of rosette flatness (as in Fig. 1), showed that SrCl_2_ shifts the 3D growth transition to lower cell numbers. In the case of the highest SrCl_2_ concentration, the transition occurred by the 5-cell stage. For both (B) and (C), results were from a total of 100 rosettes pooled from 3 experiments, with at least 8 rosettes for each size class for both SrCl_2_ concentrations. **(D)** Relative residual stress, as measured by maximum initial recoil velocity (rescaled, as in 4E, F, by the length scale of the average cell diameter of 5 μm and the time scale of the average division time of 6 hrs.) after laser ablation of single cells (as in figure 4E, F), increased in rosettes of 4-7 cells under 5mM SrCl_2_ treatment compared to untreated rosettes. These data demonstrate that increased residual stress is correlated with altered cell packing and hence, altered rosette morphology. Points represent means and error bars represent standard error of the mean from 41 total measurements pooled from two experiments, with at least 4 rosettes from each size class.

Using laser ablation experiments, we found that relative residual stress as determined by maximum initial recoil velocity (as in Fig. 4E, F) was significantly increased for SrCl_2_-treated 4-7 cell rosettes relative to untreated rosettes (Fig. 5D). The increase in residual stress in conjunction with the 2D to 3D growth transition at lower cell numbers, supported the hypothesis that ECM-constrained proliferation is a key driver of the 3D transition in rosette morphogenesis. Interestingly, for the 8-cell-stage and higher, we did not find a significant increase in residual stress in SrCl_2_-treated rosettes compared to untreated rosettes. This is consistent with cells exerting maximum growth pressure on their neighbors at the 8-cell stage and above. Taken together, these results confirm important model predictions by demonstrating that material properties of the ECM can affect morphogenesis, which highlights the central role of the ECM in sculpting rosette morphology.

### Amount of ECM, cell shape, and ECM stiffness as control parameters for morphogenesis

To formalize and test our hypothesis of morphogenesis shaped by ECM constraint (Fig. 4A, B), we next developed a cell-based computational model to simulate rosette development. Because development involves few cells (ruling out continuum modeling) in a low Reynolds number environment where inertial forces play a negligible role (49, 50) we developed particle-based simulations akin to Brownian dynamics, but neglected the role of thermal fluctuations given the large size of the cells and aggregates (51). In the model, the ECM and cells were represented by a system of interacting spherical particles (Fig. 6A). This particle representation also allowed us to capture the discrete and stochastic nature of cell division and the stochastic nature ECM secretion as well the polarity of cell division and ECM secretion. Each cell in the model was composed of three linked spheres to capture cell shape and for computational tractability, with a small sphere representing the basal pole of the cell, a larger sphere representing the cell body, and the largest representing the collar exclusion region. Cells interacted sterically with one another. The ECM was modelled as a system of small spheres with attractive interactions in order to capture the complex shapes the ECM can take on (Fig 4C) as well as its deformability. ECM particles similarly shared attractive interactions with the basal poles of cells. Cells in the model were allowed to divide stochastically, with the division plane orientation around the apico-basal axis determined by the previous division (consistent with observations of rosette development from Fig. 3 and (17)), and ECM particles were secreted stochastically at a constant rate from the basal pole of non-dividing cells (see Methods for a more detailed description of the model and simulations).

**Figure 6.**
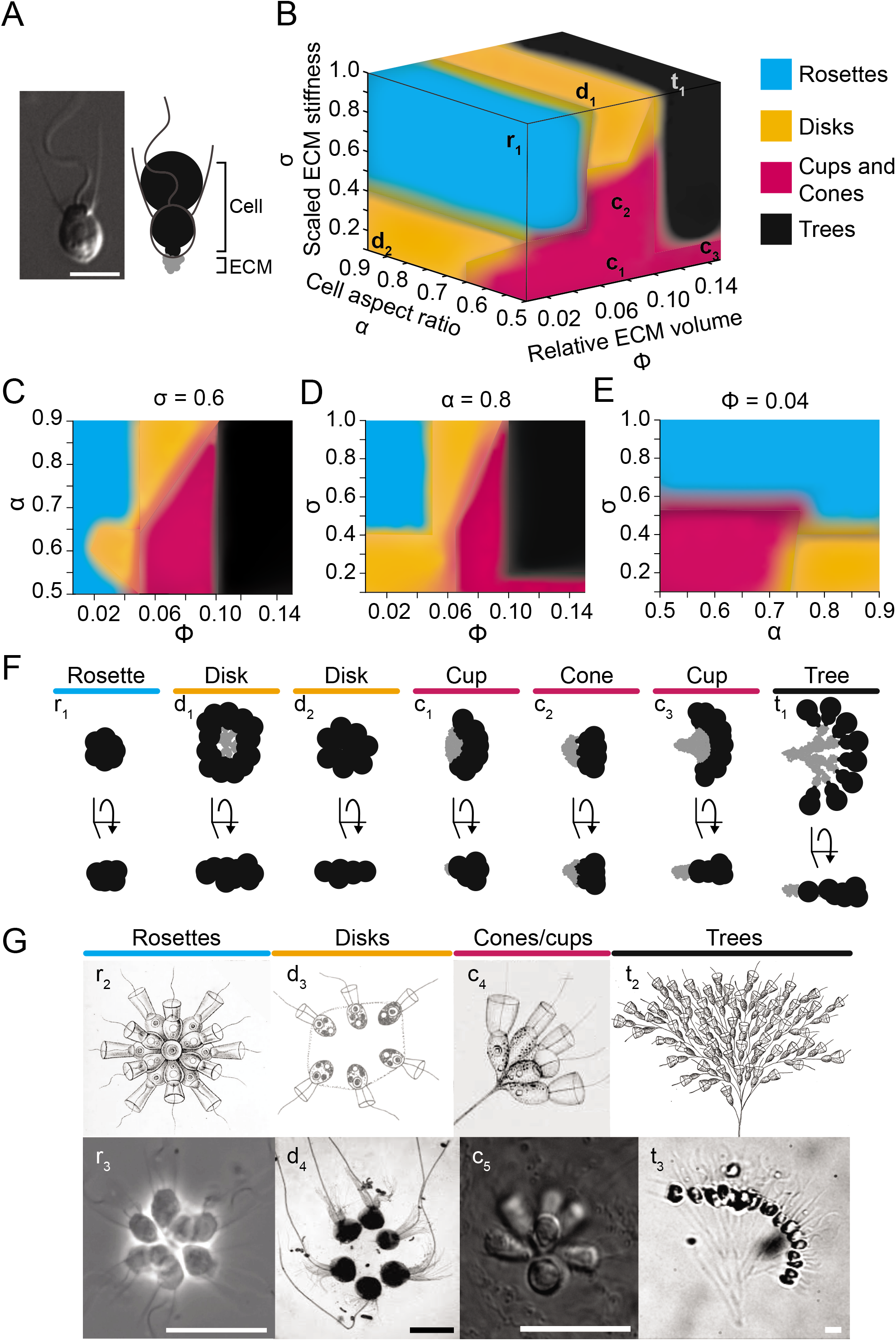
A simple model shows that amount of ECM, cell shape (aspect ratio), and ECM stiffness tune multicellular morphogenesis. The model incorporates simple cellular and physical interactions, including ECM secretion and cell division, cell-cell steric interactions, and ECM adhesion. Three main parameters describe the system: cell aspect ratio, α, scaled ECM stiffness, σ, and relative ECM volume, ϕ. **(A)** An image of a choanoflagellate (adapted from (16)) next to a simulation snapshot to illustrate how cell geometry is modeled by three linked spheres (black) and ECM is modeled by small spheres (grey) secreted at the basal pole of cells. In the model, cells interact sterically with one another, and ECM spheres have adhesive interactions with one another and with basal cell particles. Scale bar = 5 μm. **(B)** The morphospace of ECM-based colonial morphologies generated by simulations can be broken into four regions: rosettes, disks, cones/cups, and trees as denoted by colors as indicated in the legend. The lower-case letters indicate approximately the point in the morphospace occupied by the corresponding simulated colony in panel F. **(C-E)** Orthogonal planes through the displayed morphospace, with the parameter of fixed value noted above each plot, illustrate how changing two parameters while keeping the third fixed affects morphology. Colors indicate morphological classification as in panel B. **(C)** Scaled ECM stiffness is constant (σ=0.6). **(D)** Cell aspect ratio is constant (α=0.8). **(E)** Relative ECM volume is constant (ϕ=0.04). **(F)** Representative simulated colonies for each of the regions are displayed in two orthogonal views (**r_1_**=rosette with α=0.7, σ=0.8, and ϕ=0.04; **d_1_**=disk with α=0.75σ=0.85, and ϕ=0.075; **d_2_**=disk with α=0.8, σ=0.15, and ϕ=0.02; **c_1_**=cone with α=0.55, σ=0.5, and relative ECM volume=0.08; **c_2_**=cup with α=0.6, σ=0.2, and ϕ=0.09; **c_3_**=cup with α=0.9, σ=0.12, and ϕ=0.13; and **t_1_**=tree with α=0.65, σ=0.9, and ϕ=0.12). Note that d_2_ and c_3_ represent extreme ends of the morphospace to better illustrate, along with the other representative simulation snapshots, how changing the model parameters affects the simulated morphologies. **(G)** Simulated colonial morphologies are reminiscent of morphologies of colonial choanoflagellates found in nature. (**r_2_**) *Codonosiga botrytis* (86). (**r_3_**) *Salpingoeca rosetta* (87). (**d_3_**) *Proterospongia haeckelii* (88) (Ertl after Lackey). (**d_4_**) *Salpingoeca amphoridium* (10). (**c_4_**) *Codosiga umbellata* (53). (**c_5_**) Uncharacterized environmental isolate collected from Mono Lake by Daniel Richter, *Salpingoeca sp*. (**t_2_**) *Codosiga cymosa* (52) (Calkins after Kent). (**t_3_**) Uncharacterized environmental isolate from a tide pool in Curaçao, *Salpingoeca sp*. Scale bars = 10 μm.

In this simplified model of rosette morphogenesis, three main parameters characterized the system: cell aspect ratio (length along apical/basal axis vs. equatorial diameter), α, amount of ECM relative to total cell volume, ϕ, and relative stiffness of the ECM (in terms of the strength of ECM-ECM adhesion bonds relative to the force exerted by growing and dividing cells), σ. Simulations with parameter values constrained by cell and ECM morphology data collected as part of this study showed that this simple model was sufficient to recapitulate rosette morphogenesis, including the expected 3D transition at the 8-cell stage (Fig. S5). Furthermore, simulations showed that rosette morphogenesis was robust to a range of scaled ECM stiffness values (Fig. 6B-E, S5) and to the stochasticity of cell divisions (Fig. S5). We did find that simulations fail to recapitulate all aspects of rosette morphogenesis, most saliently, the growth scaling (Fig. 2D) and the absolute magnitudes of flatness and sphericity (Fig. 2E, S5). We expect, however, that a more detailed treatment of the mechanics of cells and ECM may capture these aspects of rosette morphogenesis more accurately, but such a detailed model is beyond the scope of the present study.

Exploration of the effects of different parameter values revealed that the model captures a range of different colonial morphologies (Fig. 6B, F). This space of forms and associated model parameters constitutes a theoretical morphospace (51) of ECM-based colonial choanoflagellate morphologies given this simplified model of morphogenesis. Interestingly, some of the simulated forms resembled colonies, such as tree-like structures (found in *Codosiga cymosa* (52) and an uncharacterized *Salpingoeca sp*., Fig. 6Gt_2_, t_3_) or cups (found in *Codosiga umbellata* (53) and another uncharacterized *Salpingoeca sp*., Fig. 6Gc_4_, c_5_), that have been previously reported in other choanoflagellate species (Fig. 6F, G). We found that colony morphogenesis is particularly sensitive to α and ϕ, changes in each of which can lead to dramatic changes in predicted multicellular forms. For example, holding the other two parameters fixed, increase in ϕ alone would be predicted to drive a change from rosettes to disks or cups and from cones to trees (Fig. 6C, D). Colony morphogenesis was also affected by changes in σ, but the effects tended to be subtler, such as changes in cell packing over a relatively wide range of values (Fig. S5). In contrast with changes in ϕ, increase in σ alone was either not predicted to lead to any transitions in predicted colony morphology type or, at most, lead to single transitions such as from rosettes to disks (Fig. 6D, E). These results demonstrate that basal secretion of a shared ECM constitutes a robust yet flexible mechanism for regulating multicellular morphogenesis. Furthermore, these results made specific predictions about different colony morphologies corresponding to specific cell morphologies and relative ECM volumes and stiffnesses.

## Discussion

Our quantitative analyses, experimental perturbations, and simulations allowed us to understand the process by which single cells of *S. rosetta* gives rise to multicellular rosettes. We found that the earliest stages of rosette morphogenesis proceed through 2D anisotropic growth, which is stereotypically followed by a transition to 3D isotropic growth. In particular, we found that the basal ECM secreted by cells during rosette development physically constrains proliferating cells, and thereby drives a stereotyped morphogenetic progression in the absence of strict cell lineage specification and division timing. Simulations showed that this simple mechanism, the regulated basal secretion of ECM, is sufficient to not only recapitulate rosette morphogenesis but yield a morphospace that can not only explain the multicellular morphology of *S. rosetta* but also that of other species of colonial choanoflagellates. These results emphasize the importance of the choanoflagellate ECM for morphogenesis and should encourage future studies of its composition, physical properties, and regulation.

The importance of the basal ECM revealed in this study may generalize to other choanoflagellate species and colonial morphologies. Our simulations predict that differences in ECM levels (resulting from differing rates of biosynthesis, secretion, or degradation), cell shape, and in ECM stiffness relative to cells are sufficient to explain the existence of radically different colony morphologies across diverse choanoflagellates. Measurements and comparisons of ECM levels (ϕ), cell shape (α), and ECM stiffness (σ), in diverse colonial choanoflagellates will be crucial to validate the model, and deviations from the predictions of the model could point to additional regulatory mechanisms.

From a broader perspective, rosette morphogenesis shows interesting parallels to mechanisms underlying morphogenesis in diverse other taxa. In terms of physical mechanisms, the constrained proliferation of cells that occurs during rosette development generates crowding stresses like those that regulate morphogenesis by animal epithelia (54, 55), snowflake yeast (56), and bacterial biofilms (26). In epithelia, compaction of cells due to crowding has been proposed as a general signal for cellular processes underlying tissue homeostasis such as apoptosis and extrusion (57–60). Further, accumulation of stress due to crowding of cells produces a jamming-like behavior that has been proposed as a generic constraint on the development of multicellular systems with fixed cell geometry (56). Due to the generality of physical constraints on cell packing, it is plausible that such phenomena acted both as constraints and regulatory mechanisms in the development and morphogenesis of early animals and their ancestors.

Cellular mechanisms of rosette morphogenesis are also shared with other multicellular systems. Our results demonstrate that the regulation of basal ECM sculpts the multicellular morphology of rosettes. Thus, our biophysical studies have converged on results from genetic screens in *S. rosetta* that implicated animal ECM gene homologs in the regulation of rosette development, including a C-type lectin (18) and predicted glycosyltransferases (19). The basal ECM of rosettes is reminiscent of the basal lamina, a basally secreted layer of ECM that underpins animal epithelia and regulates tissue morphogenesis by constraining cell proliferation (29) including in *Drosophila* wing and egg chamber development (54, 61, 62), branching growth during lung and salivary gland development (63, 64), notochord expansion (65), lumen elongation (66), and in tumor growth in mammary epithelia (45). The ECM also sculpts morphogenesis in *Volvox*, in which defects in ECM composition disrupt morphogenesis (67, 68), and in bacterial biofilms, in which the ECM can constrain cells and thereby drive 3D morphogenesis (26, 69). Remarkably, some bacteria form multicellular rosettes in a process that is mediated by basal ECM secretion (70, 71).

Altogether, the principles that we can glean from the simplicity of choanoflagellate morphogenesis holds the promise of revealing general principles by which biological and physical mechanisms shape morphogenesis more broadly.

## Methods

### Choanoflagellate strains and culture

Two strains of *S. rosetta* were used for the experiments in this study: one grown solely in the presence of the non-rosette inducing bacterium *Echinicola pacifica* (72), a strain called SrEpac (73) and the other grown solely in the presence of the rosette inducing bacterium *Algoriphagus machipongonensis* (74), a strain called PX1 (23, 75).

SrEpac was grown in 5% Sea Water Complete (SWC) media at 22°C. Sea Water Complete media consisted of 250 mg/L peptone, 150 mg/L yeast extract, 150μL/L glycerol in artificial sea water and was diluted to 5% by volume in artificial sea water to make 5% Sea Water Complete media. Artificial sea water (ASW) consisted of 32.9 g Tropic Marin sea salts (Wartenberg, Germany) dissolved in 1L distilled water for a final salinity of 32-27 parts per thousand. SrEpac was passaged either 1:10 into 9mL fresh 5% SWC once a day or 1:20 every other day into 9mL fresh 5% SWC to stimulate rapid proliferation and maintain log-phase growth. Cells were grown in 25cm^2^ cell culture flask (Corning).

PX1 was grown in 25% Cereal Grass media (CGM3) at 22°C. Cereal Grass media consisted of Cereal Grass (Basic Science Supplies) added to ASW at 5g/L, steeped for 3.5 hours and then filtered. This media was then diluted to 5% by volume in ASW in order to make 25% CGM3. PX1 was passaged 1:5 into 9mL of fresh 25% CGM3 every two to three days to stimulate rapid proliferation and maintain log-phase growth. Cells were grown in 25cm^2^ cell culture flask (Corning).

### Rosette induction

Rosette development from single cells was stimulated by the addition of outer membrane vesicles (OMVs) isolated from *Algoriphagus* bacteria (23) to SrEpac cultures. To isolate OMVs, *Algoriphagus* was first grown in 200mL of SWC at 30°C for 48 hrs. on a shaker. Bacterial cells were then pelleted, and the cell free supernatant was sterile filtered, then spun at 36,000 × g for three hours at 4°C (Type 45 Ti rotor, Beckman Coulter) to pellet OMVs. Finally, OMVs were resuspended in 1.5 mL ASW. To induce rosette development, OMVs were added to SrEpac at a concentration of 1:2000 by volume. This concentration led to >90% of cells in rosettes by 48 hrs. post-induction.

### Electron microscopy

*Algoriphagus* OMV induced SrEPac cultures (48 hours post-induction) were concentrated by centrifugation (1200xg for 5 min). Colonies were resuspended in 5% BSA in ASW, high pressure frozen using a Leica EM PACT2, and fixed by freeze substitution in 0.01% OsO_4_ + 0.2% uranyl acetate in acetone (76). Samples were resin embedded in Epon Araldite (Embed-812) (77), cut into 80 nm sections, and then imaged using an FEI Tecnai 12 transmission electron microscope.

### Rosette cell number quantification

Rosettes from rapidly growing PX1 cultures (*S. rosetta* co-cultured with *Algoriphagus*) were concentrated to 5x by centrifugation (1500xg for 10 min) and resuspended by vigorous pipetting in fresh 25% CGM3 media. Rosettes were then gently adhered to a poly-D-lysine coated coverslip (FluoroDish, World Precision Instruments, Inc), which had been washed three times using 25% CGM3. Rosettes were observed using a Leica DMIL microscope with a 20x objective (Leica, N Plan, 0.35 NA), and cells were manually counted. In general, rosettes were not overlapping, and cells were deemed to belong to a rosette when oriented radially outward about a central focus.

### Quantitative morphology analysis pipeline

For morphological analysis of rosettes, SrEpac cultures were first induced to form rosettes as described above. After 24 hours, developing rosettes were pelleted by centrifugation (1500xg for 10 min) and resuspended in fresh ASW by vigorous pipetting in order to minimize bacteria and to break apart any chains that might be mistaken for rosettes. Rosettes were then deposited onto a poly-D-lysine coated coverslip (FluoroDish, World Precision Instruments, Inc), which had been washed three times using ASW. Rosettes were stained by overloading with LysoTracker Red DND-99 (ThermoFisher Scientific) at 1:200 dilution, which reliably stains the entire cell body. Next, z-stack images of stained rosettes were acquired on a Zeiss 880 laser scanning confocal microscope using a 40x water immersion objective (Zeiss, C-Apochromat, 1.2 NA) and illumination with a 561 nm laser (Zeiss). Importantly, pure water, and not water immersion oil, was used to minimize coverslip deflection during imaging.

After image acquisition, z-stacks were registered using the Stackreg plugin in FIJI (78, 79). Aligned z-stacks were deconvolved using the Parallel Iterative Deconvolution v1.12 plugin in FIJI (78, 79). For deconvolution, the Wiener Filter Preconditioned Landweber method (WPL) with stock settings and a theoretical pointspread function for the imaging system generated using the Diffraction PSF 3D plugin in FIJI (78, 79) were used. Aligned, deconvolved z-stacks were then segmented using Imaris v3.8 (Bitplane, Belfast). First, the images were median filtered with a 3×3×1 kernel and smoothed using a Gaussian filter with a sigma of 0.24 microns. Intensity thresholds for local intensity segmentation and thresholds for size and shape filters to exclude extraneous objects such as bacteria within the analysis region were then chosen based on segmentation of a few rosettes from each sample and then kept the same for all rosettes in the sample. Individual cells in rosettes were segmented using the Split Touching Objects option. Segmentation of each rosette was manually inspected, and any improperly segmented cells were manually split and fused as necessary. Statistics of segmented rosettes, including number of cells, cell positions, orientations, sizes, and shapes were exported to MATLAB release 2016a (Mathworks, Natick) for additional morphological analysis.

Rosette volume was measured by determining the convex hull of cell positions. Maximum rosette width was measured by the maximum distance between cells in rosettes. To further evaluate rosette morphology, principle axes of rosettes were determined by principle components analysis of cell positions. Flatness (*F*) and Sphericity (*S*) of rosettes were computed from these principle components where *F* = 1 − *C*/*B*, and 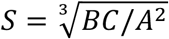 where *A, B*, and *C* are the principle axes in descending order of magnitude. The packing of cells was then quantified by the average number of neighboring cells over all cells in rosettes as determined by a Voronoi tessellation (80) of cell positions. Finally, rosettes were then binned by cell number for the final analysis of morphological progression.

### Cell lineage analysis

Rapidly growing SrEpac cultures were induced to form rosettes, and 9 hours post induction, induced cells were concentrated to 5x by centrifugation and resuspension in 5% SWC (1200xg for 5 min, initial volume 15mL resuspended in 3mL) and then deposited in a 200 μL droplet on a poly-D-lysine coated coverslip (FluoroDish, World Precision Instruments, Inc). Cells were imaged in phase contrast or DIC on either a Zeiss Axio Observer Z1 with a 20x (Zeiss, Plan-Apochromat, 0.8 NA) objective or a Leica DMI6000B with a 20x (Leica, Plan-Apochromat, 0.7 NA) objective at 1 frame/minute for 16 hours. Cell positions were tracked using the Manual Tracking plugin in FIJI (78, 79), and division events were tracked and recorded manually. For analysis of chain cell lineages, the cells were not induced to form rosettes, but otherwise all previous steps were followed.

### ECM measurements

Rosettes were prepared as in the “Quantitative morphology analysis pipeline” section (QMAP). Additionally, to label the ECM, fluorescein labeled Jacalin (Vector Labs, FL-1151) at a 1:400 dilution was added to the concentrated rosettes. Imaging also followed the QMAP with additional sequential illumination with a 488 nm laser to excite the fluorescein. Z-stack images were processed and analyzed following QMAP with the exception of post-processing in MATLAB, as cell and ECM volumes were exported directly from Imaris.

### Laser ablation

For laser ablation, an upright Olympus BX51WI microscope (Olympus Corporation) equipped with Swept Field Confocal Technology (Bruker) and a Ti:Sapphire 2-photon Chameleon Ultra II laser (Coherent) were used. The 2-photon laser was set to 770 nm and ablation was performed using three 20 ms pulses. A 60x water dipping objective (Olympus, LUMPlanFLN, 1.0 NA) was used for imaging. Images were captured using an EM-CCD camera (Photometrics). The following emission filter was used: Quad FF-01-446/523/600/677-25 (Semrock). PrairieView Software (v. 5.3 U3, Bruker) was used to acquire images.

Rosettes were gently adhered to a coverslip using poly-D-lysine and stained with lysotracker (as described above). Individual cells in rosettes were ablated, and the subsequent recoil, which is proportional to the elastic stress (7, 39, 43), was recorded at a frame rate of 1/0.48 s. Images were registered using the StackReg plugin in FIJI (78, 79) to correct for small movements of the rosette colony due to flagellar motion during acquisition of images. Recoil velocities were measured in the frames following ablation by particle image velocimetry (PIV) using PIVlab software in MATLAB (81). Settings for PIV included four direct Fourier transform correlation passes with window sizes of 64, 32, 16 and 8 pixels and corresponding step sizes of 32, 16, 8, and 4 pixels. To reject noise and erroneous velocities, filters of 7 standard deviations about the mean and local median filters with a threshold of 5 and epsilon of 0.1 were applied. Finally, any remaining velocity measurements not corresponding to displacements of cells in rosettes were manually rejected. Recoil velocities were measured in the subsequent 3 frames following ablation by radial scans about the circumference of the rosette, and the maximum measured velocity was selected.

### Strontium treatment

For strontium treatment of rosettes, SrEpac cultures were centrifuged at 1500g for 10 to pellet all cells and resuspended in 5% SWC media containing added SrCl_2_ to a final concentration of either 0, 2.5, or 5 mM. These cells were then induced to form rosettes as described above. Morphological analysis and laser ablation were also conducted as described above.

For cell growth assays (Fig. S4), SrEpac cultures were prepared as described in the preceding paragraph but were not induced to form rosettes. Cells were then plated into 12-well plates (Falcon) at an initial density of 20000 cells/mL. To determine cell density, cells were counted using a hemocytometer (Hausser Scientific) viewed in phase contrast on a Leica DMIL microscope with a 20x (Leica, N Plan, 0.35 NA) objective. Cells were counted at 4, 24, and 28 hours. Growth rates were then determined by exponential fits to the log-phase of growth obtained using the Curve Fitting application in MATLAB.

### Simulations

Cells and ECM were modelled as spherical particles (Fig. 6) with interactions that allowed us to tune the various morphological and material properties we wished to investigate. The particle representation allowed us to capture both the relevant geometric aspects of colony formation including polarized cell divisions and ECM secretion as well as the discrete and stochastic nature of these processes.

### Cells

Each cell was composed of three linked particles with diameters *d*_1_, *d*_2_, and *d*_3_ representing the basal pole, cell body, and collar and in ascending order of magnitude, to capture cell geometry. Cell particles interacted sterically with one another via the hard-sphere Weeks-Chandler-Andersen (WCA) potential (82):

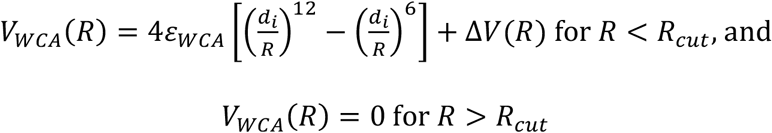

where *R* is the interparticle distance; *d_i_* with *i* = 1, 2, 3 is the cell particle diameter; *ε_WCA_* sets the force of repulsion upon overlap; Δ*V*(*R*) = *V_LJ_*(*R_cut_*) where *V_LJ_* is the Lennard-Jones potential (82):

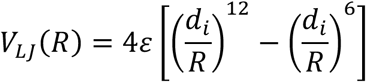

with *ε* = *ε_WCA_*; and 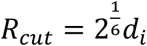 is the cutoff distance for the potential set to the diameter of the cell particle. For a given cell, cohesion of particles was maintained using a finitely extensible nonlinear elastic (FENE) potential:

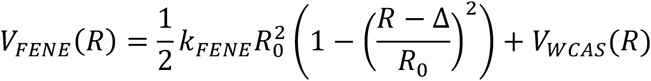

where *k_FENE_* sets the strength of the potential; *R*_0_ = *d*_1_ (set to maintain cohesion); *R* is the interparticle distance; 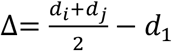 where *d_k_* is the diameter of the *k^th^* particle; and

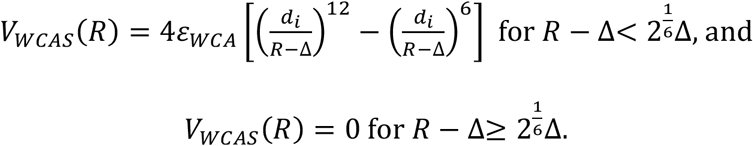

Additionally, a harmonic potential acting between the basal and apical cell particle was used to keep cells straight and elongated:

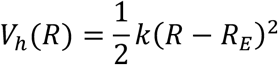

where the spring constant *k* = *k_h_* sets the strength of the potential, and rest length *R_E_* = 2(*d*_1_/2 + *d*_2_ + *d*_3_/2) was chosen to be large enough to ensure cell elongation. For simplicity, the mass of all cell particles was the same, and the friction coefficient was otherwise determined by viscosity, *η* and the particle diameter, *d*: *γ* = 6*πηd*.

### ECM

ECM was composed of small particles with diameter *d_ECM_* ≪ *d*_1_. To maintain ECM cohesion (while preventing divergence in energy) and volume, allow for ECM deformations and shape transformation, and for computational tractability, ECM-ECM particle interactions were also modeled by a modified Lennard-Jones potential (83), which reduces interparticle repulsion and better describes a condensed state, such as ECM:

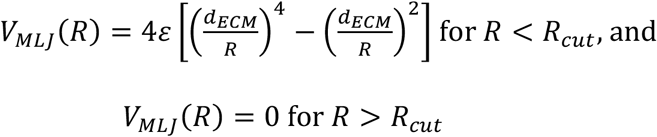

where *R* is the interparticle distance, *ε* = *ε_ECM_* sets the strength of ECM adhesion, and 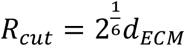 is the cutoff distance for the potential.

### Cell-ECM interactions

To capture adhesive interactions while preventing particle overlap and for computational tractability, cell-ECM adhesion was modeled with a modified Lennard-Jones potential between basal cell particles and ECM particles:

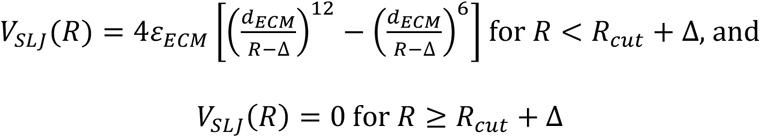

where 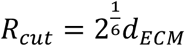, and 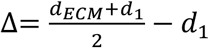.

### Cell division

Cells were allowed to divide stochastically, with probability 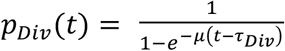 where *τ_Div_* is the cell cycle time, and 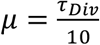 sets the variability in division timing. The division plane orientation around the apico-basal axis was set by the previous division. During division, cell particles are replicated and shifted slightly (starting with a separation 0.25*d_i_* for particle *i*) in the direction perpendicular to the division plane by an offset *D_Div_*. Dividing cell particles then push one another apart under the influence of a harmonic potential with increasing rest length: 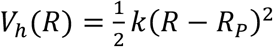 where the spring constant *k* = *k_Div_* sets the strength of the potential, and hence, how much force the cells can exert during growth and division; and rest length *R_P_* increases with each timestep in the simulation by an amount 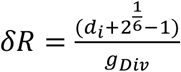 where *g_DIV_* is the number of timesteps over which division occurs for particle *i*. Cell division is complete once *R_P_* ≥ *d_i_* for all particles. With this implementation of cell division, we approximated both cell growth (while particles are still overlapping) and division.

### ECM secretion

ECM particles were secreted stochastically from non-dividing cells at a constant rate with a probability 1 − *p_Div_*(*t*) from the basal pole of non-dividing cells. Secretion always occurred in the direction of the apico-basal axis of cells. Similarly to division, ECM particles were extended from the basal pole according to a harmonic potential with *k* = *k_ECM_* setting the force with which ECM particles were secreted; the rest length *R_P_* increased with every time step by 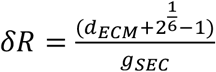 where *g_SEC_* is the number of timesteps over which secretion occurs. Secretion is complete once 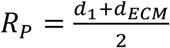.

### Running simulations

Simulations were carried out using Fortran and followed Brownian dynamics (84):

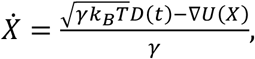

where *X* and *Ẋ* are position and velocity; *U*(*X*) is the sum of all interaction potentials acting on a given element of the system (particle), so −∇*U*(*X*) with ∇ as the gradient operator, is the force resulting from the total interaction potential on a given element of the system (particle); *γ* is the friction coefficient, *k_B_* is Boltzmann’s constant, *T* is temperature, and *D*(*t*) is a delta correlated, stationary Gaussian process with 0 mean. For simplicity, the mass of all particles was the same, and the friction coefficient was otherwise determined by viscosity, *η* and the particle diameter, *d*: *γ* = 6*πηd*. A Verlet integration algorithm was used to update the positions of the spheres at each timestep in the simulation (85). Code for running simulations is available on GitHub: https://github.com/truizherrero/choanoflagellate_colonies.

### Simulation analysis

In the model, three main parameters corresponding to physical aspects of choanoflagellate cells and rosettes describe the system: 1) cell aspect ratio, defined to be the length to width ratio of the three particle system: 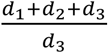; 2) scaled ECM stiffness relative to the force exerted during cell growth and division: 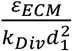; and 3) The ECM volume relative to cell volume secreted by a cell between divisions: 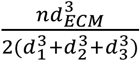 where 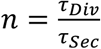 with 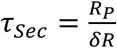 is the average number of ECM particles secreted between divisions.

Natural units for the system are the length *d*_1_, time *τ_Div_*, and energy *k_B_T*. For all simulations, the following values in system units were held fixed: 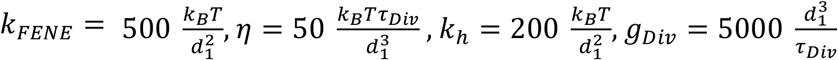, and 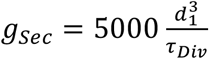. All other parameters not already fixed in the previous paragraphs were varied to explore the morphospace (Fig. 6B), with axes corresponding to the three main parameters detailed in the previous section. The timestep for simulations was 0. 001*τ_Div_*. Simulation snapshots for Fig. 6 were rendered for the visual inspection of morphologies using Python. Rosettes were defined as structures with cells completely surrounding a central region of ECM; disks were defined as structures that maintained a closed ring of cells pointing radially outward along the colony circumference with an open central region of ECM; cups and cones were defined as structures with cells clustered together, oriented in roughly the same direction, opposed to an open ECM emanating away from the basal pole of all cells; and trees were defined as structures with cells oriented in a similar fashion to those in cones but with the ECM displaying a dichotomous branching structure. Quantitative analyses of simulation results (Fig. S5) were carried out in MATLAB.

## Acknowledgements

We thank Kent McDonald and Reena Zalpuri of the Electron Microscopy Laboratory at UC Berkeley for assistance with TEM sample preparation and imaging. We would also like to thank Mary West of the CIRM/QB3 Shared Stem Cell Facility at UC Berkeley. Additionally, we thank George Oster and Danny Wells for helpful discussions concerning conceptual development of this project. Finally, we thank the members of the King Lab for critical feedback and stimulating discussions. This work was performed in part in the SSCF, which provided the Olympus BX51WI microscope with Swept Field Confocal technology. This material is based upon work supported by the National Science Foundation Graduate Research Fellowship under Grant No. DGE 1106400 (to B.T.L.), the Simons Foundation (T.R.H.) the Howard Hughes Medical Institute (N.K.), and the National Institutes of Health (NIH) under award numbers F31GM119329 (to S.L.), R01GM122375 (to S.K.), and R21EB016359 (to S.K.).

