## Supplementary Figures for "Biophysical principles of choanoflagellate self-organization"

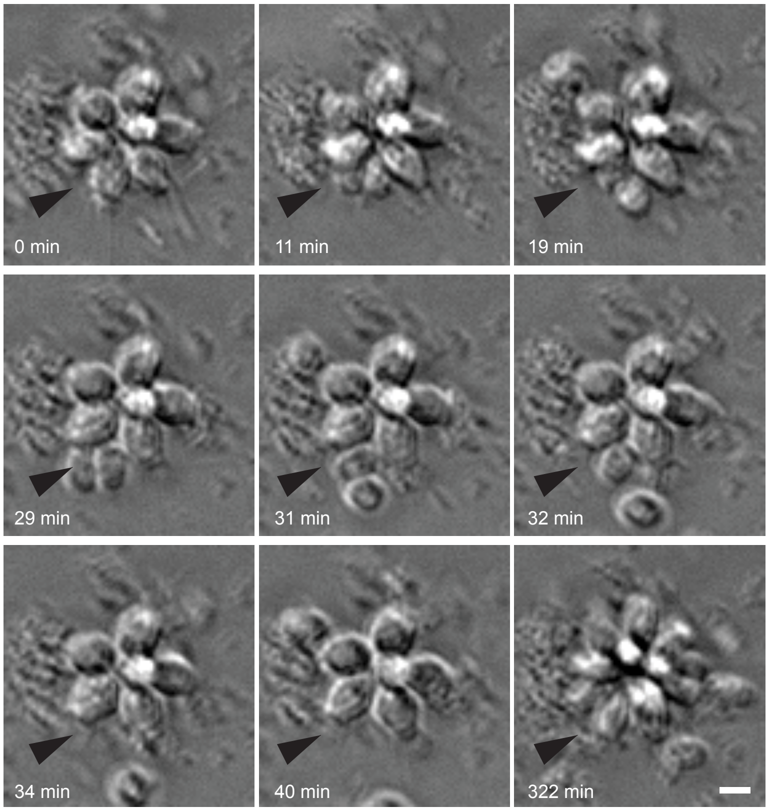

0 min
11 min
19 min
29 min
31 min
32 min
34 min
40 min
322 min

**Figure S1. Rosettes can decrease in cell number by extrusion of single cells.** Snapshots from a time series collected by DIC imaging illustrating the process by which cells are extruded from a developing rosette and swim away as solitary cells. Black arrowhead indicates location of extrusion event and white arrowheads indicate extruded cells. Scale bar = 5 μm.

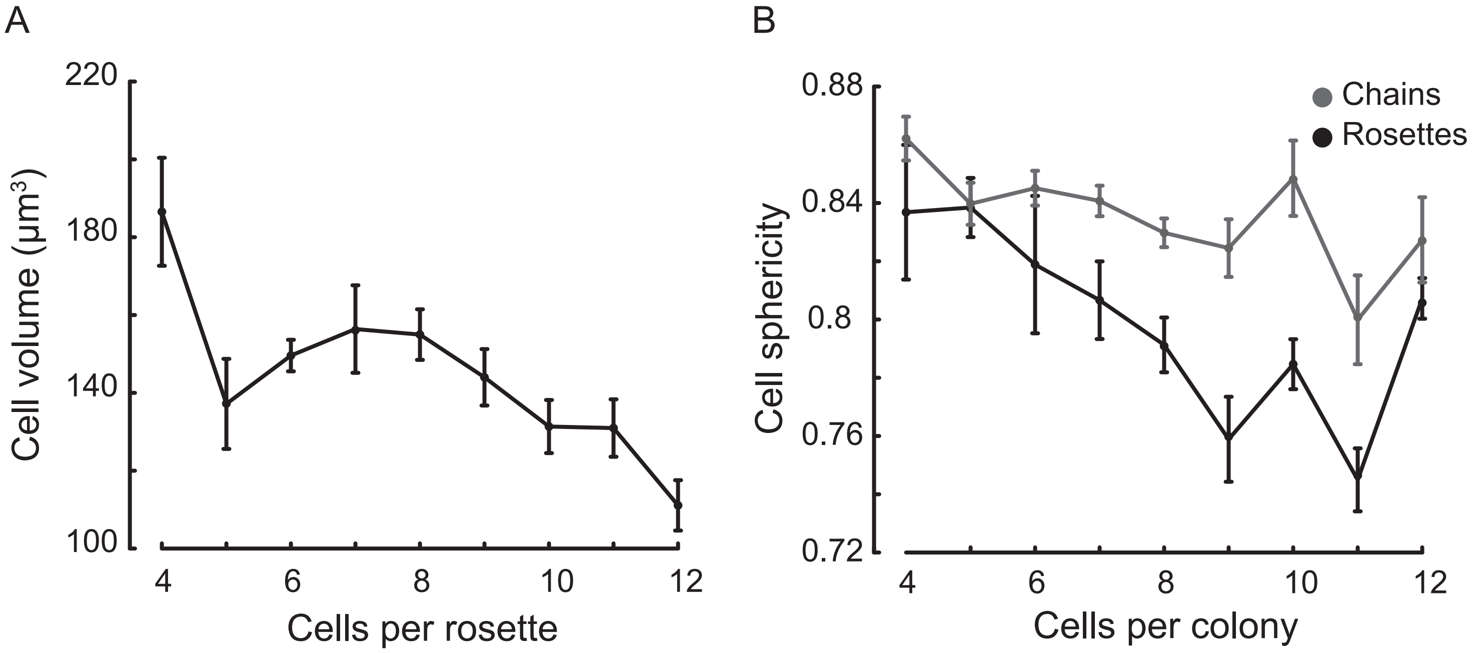

A
220
180
140
100
Cell volume (μm³)
4
6
8
10
12
Cells per rosette
B
0.88
0.84
0.8
0.76
0.72
Cell sphericity
Chains
Rosettes
4
6
8
10
12
Cells per colony

**Figure S2. Cell size and shape changes during colony development.** (**A**) Average size of cells in rosettes decreases significantly from the 4 to 5-cell stage and then experience a modest, progressive decrease in cell size after the 8-cell stage. Error bars are standard error of the mean. (**B**) Sphericity of cells as measured by $\pi^{\frac{1}{3}}(6V)^{\frac{2}{3}}/A$ (89), where V is cell volume and A is cell area, decreases in rosettes while remaining constant in chains. This is consistent with cells in rosettes either actively changing shape during development, becoming deformed by compressing due to cell packing, or a combination of the two. Data is pooled from 100 rosettes and 110 chains, with at least 8 colonies from each size class. Error bars are standard error of the mean.

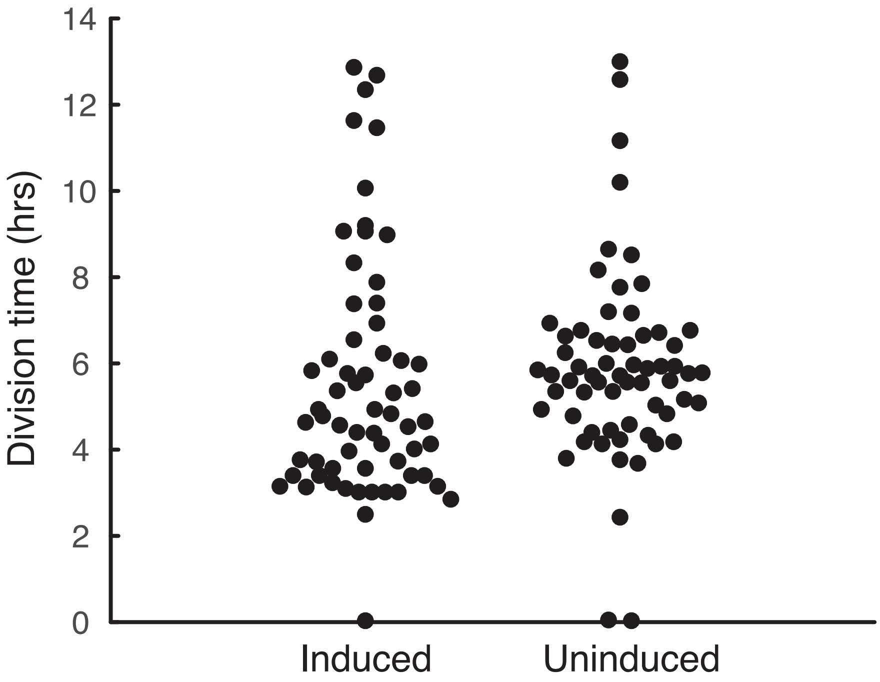

Division time (hrs)
14
12
10
8
6
4
2
0
Induced
Uninduced

**Figure S3. Division times in developing rosettes, induced cells, are slightly but significantly different from those of uninduced cells** ($p=0.03$ by Wilcoxon rank sum method). While the range of division times is the same, induced cells show a small increase in division rate (mean division time of 5.5 hrs for induced cells vs. 5.9 hrs for uninduced cells). Data is pooled from 20 different colonies for both the induced and uninduced conditions.

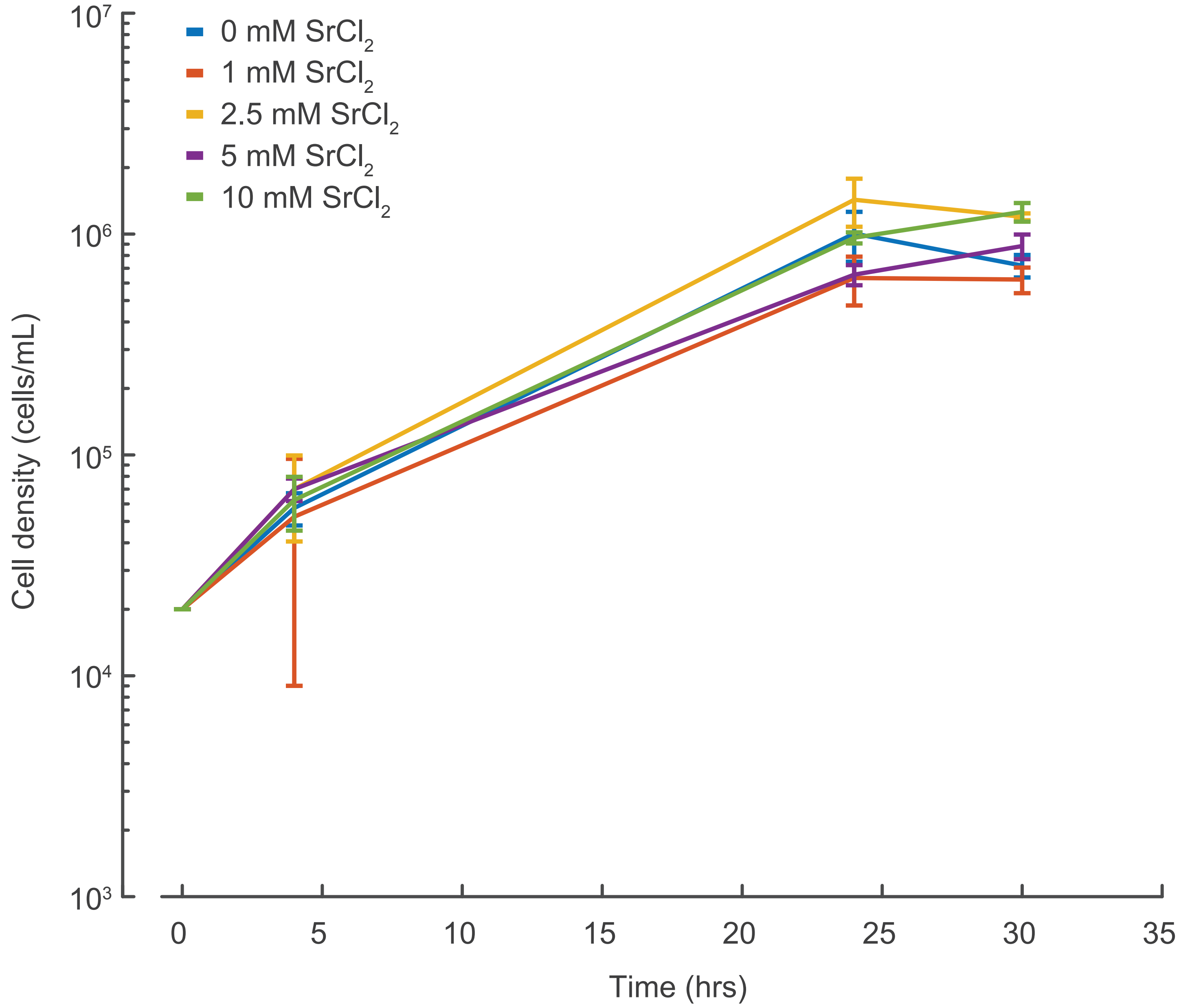


| $SrCl_2$ Concentration (mM) | Growth Rate (division/hr) |
|---|---|
| 0 | 0.15 ± 0.02 |
| 1 | 0.13 ± 0.04 |
| 2.5 | 0.16 ± 0.01 |
| 5 | 0.12 ± 0.01 |
| 10 | 0.14 ± 0.02 |

**Figure S4. $SrCl_2$ does not affect cell growth rates at up to twice the concentration used in the experiments in this study** (Fig. 5). The plot displays growth curves on a log scale for various concentrations of $SrCl_2$ and the bottom table displays corresponding growth rates derived from exponential fits of the log-phase of each growth curve.

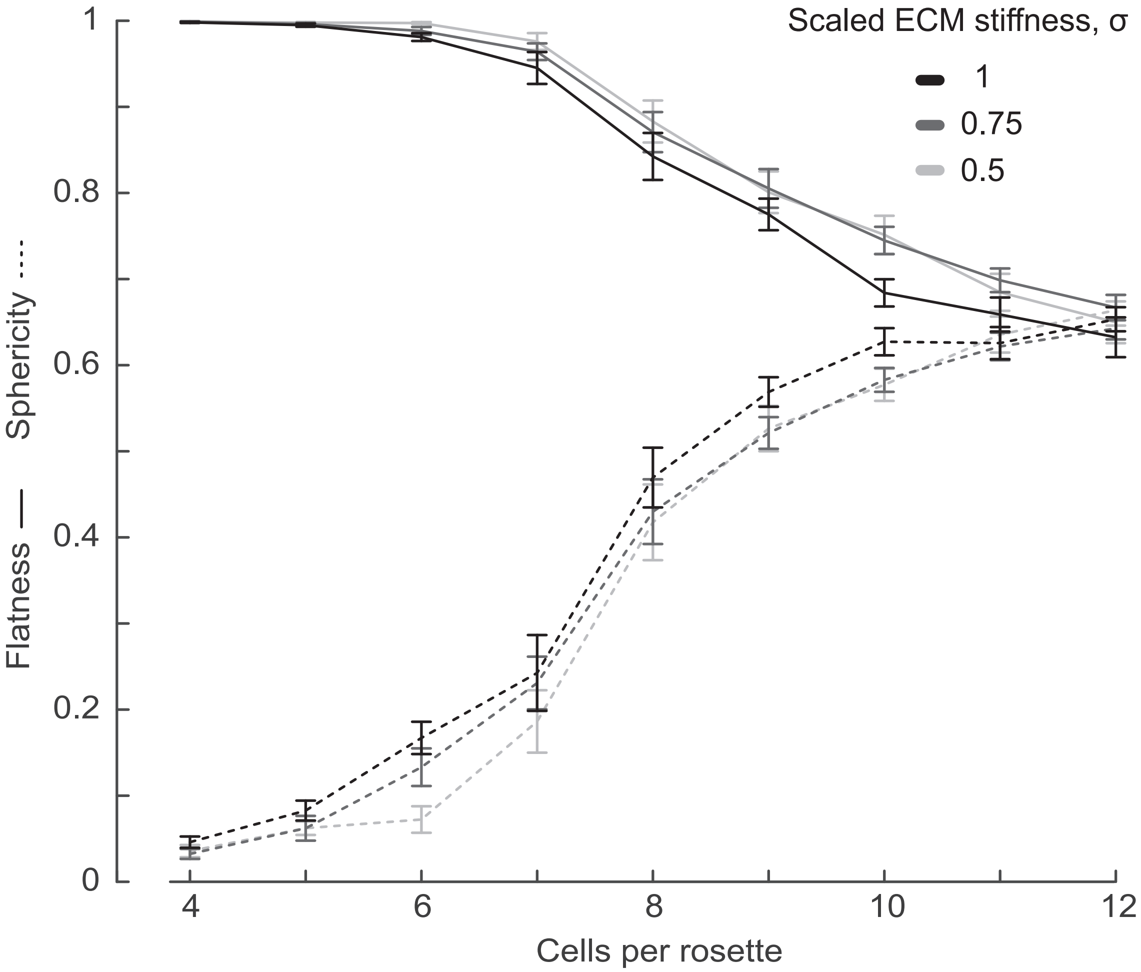

Scaled ECM stiffness, σ
1
0.75
0.5
Flatness —— Sphericity ----
1
0.8
0.6
0.4
0.2
0
4
6
8
10
12
Cells per rosette

**Figure S5. Simulations of colony development with parameters constrained by measurements of rosette and ECM morphology recapitulate rosette morphogenesis including the 3D transition at the 8-cell stage.** Simulated rosette morphogenesis is robust to the stochasticity of cell divisions and to a range of different values of scaled ECM stiffness, $\sigma$, and rosettes are increasingly 3D at lower cell numbers with increasing $\sigma$, consistent with results from experiments (Fig. 5C). The mean of 10 different simulation runs for each $\sigma$ value is plotted here, and error bars are the standard error of the mean.
